# Cell Cycle Constraints and Environmental Control of Local DNA Hypomethylation in α-Proteobacteria

**DOI:** 10.1101/091785

**Authors:** Silvia Ardissone, Peter Redder, Giancarlo Russo, Antonio Frandi, Coralie Fumeaux, Andrea Patrignani, Ralph Schlapbach, Laurent Falquet, Patrick H. Viollier

**Author notes:** Current address: LMGM, Centre de Biologie Integrative, Universite Paul Sabatier, Toulouse, France. Current address: Department of Fundamental Microbiology, University of Lausanne, Lausanne, Switzerland (Antonio Frandi). Current address: Department of Microbiology and Immunology, Harvard Medical School, Boston, MA, 02115, USA. Corresponding authors (SA), (PV).

## Abstract

Heritable DNA methylation imprints are ubiquitous and underlie genetic variability from bacteria to humans. In microbial genomes, DNA methylation has been implicated in gene transcription, DNA replication and repair, nucleoid segregation, transposition and virulence of pathogenic strains. Despite the importance of local (hypo)methylation at specific loci, how and when these patterns are established during the cell cycle remains poorly characterized. Taking advantage of the small genomes and the synchronizability of α-proteobacteria, we discovered that conserved determinants of the cell cycle transcriptional circuitry establish specific hypomethylation patterns in the cell cycle model system *Caulobacter crescentus.* We used genome-wide methyl-N6-adenine (m6A-) analyses by restriction-enzyme-cleavage sequencing (REC-Seq) and single-molecule real-time (SMRT) sequencing to show that MucR, a transcriptional regulator that represses virulence and cell cycle genes in S-phase but no longer in G1-phase, occludes 5’-GANTC-3’ sequence motifs that are methylated by he DNA adenine methyltransferase CcrM. Constitutive expression of CcrM or heterologous methylases in at least two different α-proteobacteria homogenizes m6A patterns even when MucR is present and affects promoter activity. Environmental stress (phosphate limitation) can override and reconfigure local hypomethylation patterns imposed by the cell cycle circuitry that dictate when and where local hypomethylation is instated.

**Author Summary:** DNA methylation is the post-replicative addition of a methyl group to a base by a methyltransferase that recognise a specific sequence, and represents an epigenetic regulatory mechanism in both eukaryotes and prokaryotes. In microbial genomes, DNA methylation has been implicated in gene transcription, DNA replication and repair, nucleoid segregation, transposition and virulence of pathogenic strains. CcrM is a conserved, cell cycle regulated adenine methyltransferase that methylates GANTC sites in α-proteobacteria. N^6^-methyl-adenine (m6A) patterns generated by CcrM can change the affinity of a given DNA-binding protein for its target sequence, and therefore affect gene expression. Here, we combine restriction enzyme cleavage-deep sequencing (REC-Seq) with SMRT sequencing to identify hypomethylated 5’-GANTC-3’ (GANTCs) in α-proteobacterial genomes instated by conserved cell cycle factors. By comparing SMRT and REC-Seq data with chromatin immunoprecipitation-deep sequencing data (ChIP-Seq) we show that a conserved transcriptional regulator, MucR, induces local hypomethylation patterns by occluding GANTCs to the CcrM methylase and we provide evidence that this competition occurs during S-phase, but not in G1-phase cells. Furthermore, we find that environmental signals (such as phosphate depletion) are superimposed to the cell cycle control mechanism and can override the specific hypomethylation pattern imposed by the cell cycle transcriptional circuitry.

## Introduction

DNA methylation is a conserved epigenetic modification that occurs from bacteria to humans and is implicated in control of transcription, DNA replication/repair, innate immunity and pathogenesis [1, 2]. Originally described as a mechanism that protects bacteria from invading foreign (viral) DNA [3], methyl-N6-adenine (m6A) modifications are thought to direct infrequent and stochastic phenotypic heterogeneity in bacterial cells [4, 5] and were recently implicated in transcriptional control of lower eukaryotic genomes [4, 5] and silencing in mouse embryonic stem cells [6–8].

How local changes in methylation are instated during the cell cycle remains poorly explored, even in γ-proteobacteria such as *Escherichia coli* and *Salmonella enterica*, as cell cycle studies on cell populations are cumbersome and require genetic manipulation [9]. Moreover, the replication regulator SeqA that controls the methylation state by preferentially binding hemi-methylated sequences is only encoded in γ-proteobacteria, suggesting that other mechanisms are likely operational in other systems [9, 10]. Model systems in which cell populations can be synchronized without genetic intervention are best suited to illuminate the interplay between methylation and cell cycle [11, 12]. The fresh-water bacterium *Caulobacter crescentus* and more recently the plant symbiont *Sinorhizobium meliloti* that reside in distinct environmental niches are such cell cycle model systems [13]. Akin to other α-proteobacteria, *C. crescentus* and *S. meliloti* divide asymmetrically into a smaller G1-phase cell and a larger S-phase cell and use conserved transcriptional regulators arranged in modules to coordinate transcription with cell cycle progression [13–16] (Fig 1A). In *C. crescentus*, MucR1 and MucR2 were recently shown to negatively regulate numerous promoters that are activated by the cell cycle transcriptional regulator A (CtrA) in G1-phase. MucR orthologs control virulence functions in α-proteobacterial pathogens and symbionts, but can also control cell cycle-regulated promoters in *C. crescentus* [17–20]. MucR1/2 target promoters by way of an ancestral zinc finger-like fold and both proteins are present throughout the *C. crescentus* cell cycle [17, 21, 22] (Fig 1A). By contrast, the OmpR-like DNA-binding response regulator CtrA is activated by phosphorylation and is only present in G1 and late S-phase cells [23, 24], but not in early S-phase cells (Fig 1A). The promoter controlling expression of the conserved DNA methyltransferase CcrM is among the targets activated by phosphorylated CtrA (CtrA~P) in late S-phase [15, 17, 25–27]. CcrM introduces m6A marks at sites harbouring the recognition sequence 5’-GANTC-3’ (henceforth GANTCs) once passage of the DNA replication fork leaves GANTCs hemi-methylated (Fig 1B). CcrM is an unstable protein degraded by the ATP-dependent protease Lon throughout the cell cycle [28, 29]. Since the *ccrM* gene is expressed only in late S-phase cells, the time of expression dictates when the unstable CcrM protein is present during the cell cycle. CcrM no longer cycles when it is expressed from a constitutive promoter in otherwise *WT* cells or when Lon is inactivated [28, 30].

**Fig 1.**
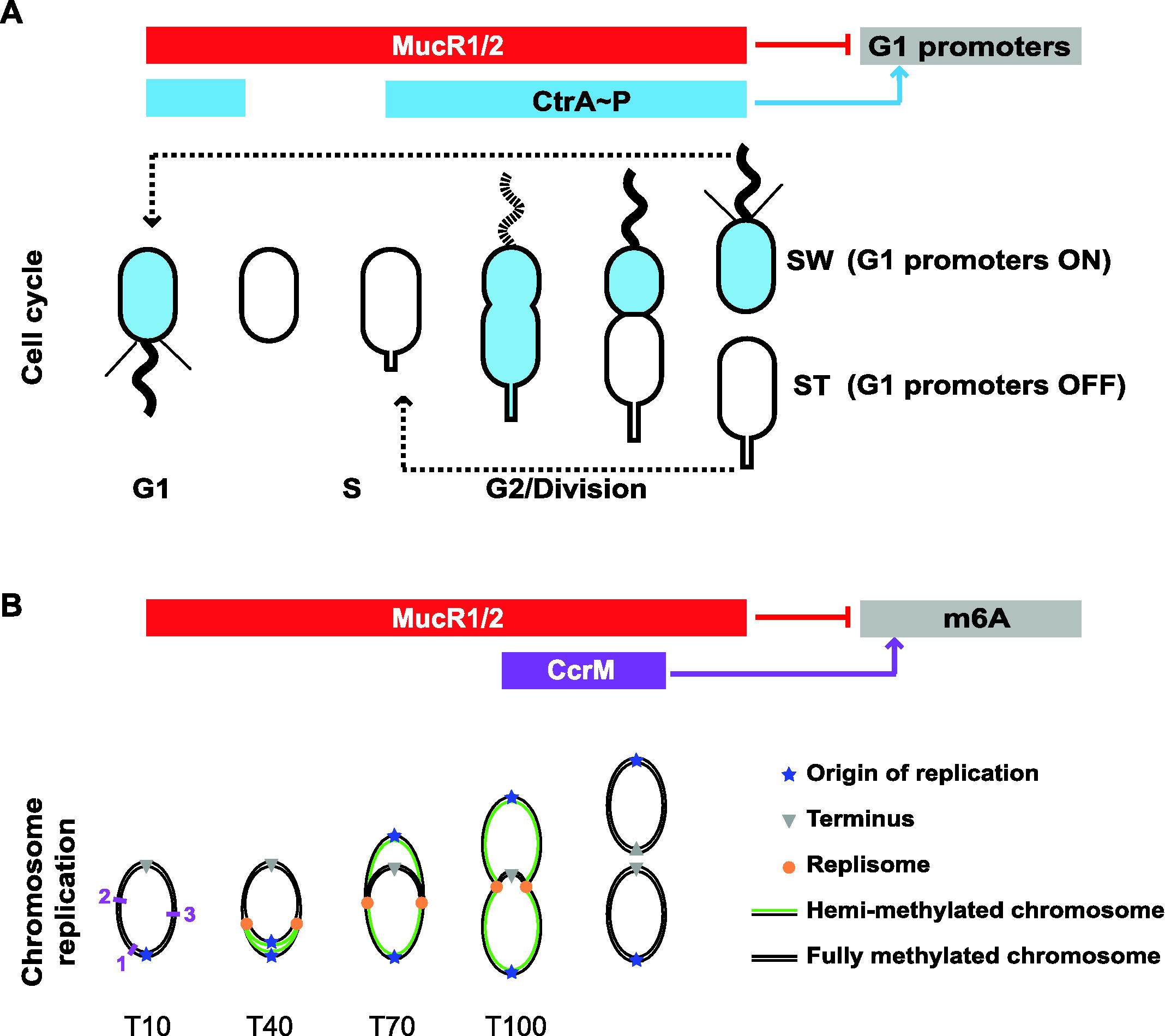
Regulation of *C. crescentus* cell cycle and methylation of the chromosome. (A) Schematic of the *C. crescentus* cell cycle and the regulatory interactions that control G1-phase promoters. Transcription from G1-phase promoters is activated by phosphorylated CtrA (CtrA~P, in blue) and repressed by MucR1/2. CtrA~P accumulates in pre-divisional and swarmer (SW) cells, and is eliminated by regulated proteolysis in the stalked (ST) cell upon compartmentalization and at the swarmer to stalked cell differentiation. The red bar indicates that MucR1/2 proteins are present at all stages of the cell cycle. (B) Schematic of the chromosome replication and adenine methylation (m6A) during the *C. crescentus* cell cycle. Chromosome replication and methylation are shown for the same cell cycle stages depicted in panel A. In non-replicative SW cells (T10) the chromosome is methylated on both strands. Upon differentiation into ST cells (T40), chromosome replication start from the origin (blue star) located at the old cell pole; progression of the replisomes (orange dots) generates hemi-methylated chromosomes (black and green lines represent respectively the methylated and unmethylated strand; T70 and T100). The cell cycle-regulated adenine methyltransferase CcrM is synthesized only in late pre-divisional cells, where it methylates the newly synthesized chromosome strands. CcrM is then specifically proteolyzed by the Lon protease. The purple numbers indicate the position along the chromosome of the *P169* (1), *P1149* (2) and *P2901* (3) promoters. T10, T40, T70 and T100 indicate the time after synchronization at which the samples for anti-MucR1 ChIP-Exo were taken (see text).

With the advent of SMRT (single-molecule real-time) sequencing it is now possible to obtain m6A-methylome information of bacterial genomes at single base pair resolution [31, 32]. A recent cell cycle methylome analysis of *C. crescentus* by SMRT-sequencing revealed the large majority of GANTCs switch from hemi-methylated to a full methylated state (m6A-marked GANTCs on both strands) at the onset of CcrM expression [12]. Interestingly, a few sites were consistently hypomethylated, indicating that site-specific mechanisms control local hypomethylation patterns. Local hypomethylation patterns may arise if specific DNA-binding proteins and/or restricted local chromosome topology block access of CcrM to such GANTCs. Here, we combine restriction enzyme cleavage-deep sequencing (REC-Seq) with SMRT sequencing to unearth hypomethylated GANTCs in the genomes of wild type (*WT*) and mutant *C. crescentus* and *S. meliloti*. We show that the conserved transcriptional regulator MucR induces local m6A-hypomethylation by preventing CcrM from accessing GANTCs during S-phase, but only when CcrM cycles. Since repression of MucR target promoters is normally overcome in G1-phase, our data suggest that MucR is unable to shield GANTCs when CcrM is artificially present in G1 cells. Lastly, we discovered that phosphate starvation promotes methylation of specific MucR-shielded GANTCs, revealing an environmental override of the control system that normally instates local hypomethylation patterns during the cell cycle.

## Results

### Identification and analysis of hypomethylated GANTCs by restriction enzyme cleavage (REC-Seq)

Detection of hypomethylated sites by SMRT-sequencing requires sufficient sequencing depth and sophisticated bioinformatic analysis to differentiate unmethylated GANTCs from methylated ones. Since unmethylated GANTCs can be conveniently enriched for in *C. crescentus* by restriction enzyme cleavage using the *Hin*fI restriction enzyme (which only cleaves unmethylated GANTCs) [33], we sought to apply *Hin*fI-based cleavage followed by Illumina-based deep-sequencing (REC-Seq) to identify hypomethylated GANTCs, similar to a previous procedure used for analysis of hypomethylated m6A sites in the unrelated γ-proteobacterium *Vibrio cholerae* [34]. We tested REC-Seq on *Hin*fI-treated genomic DNA (gDNA) from *C. crescentus* and, following bioinformatic filtering, obtained a list of unprotected GANTCs scaling with *Hin*fI cleavage efficiency (“score” in S1 Table). Since nearly all GANTCs suggested to be consistently unmethylated by SMRT sequencing [12] are represented as high scoring GANTCs in the REC-Seq (note the medium differences or limited SMRT sequencing depth may explain the differences), we concluded that REC-Seq captures hypomethylated GANTCs in scaling manner (see below where selected sites cleaved in *WT* are no longer cleaved in the Δ*mucRl/2* mutant). Since CcrM also methylates GANTCs in other α-proteobacteria [35, 36], we also determined the hypomethylated GANTCs on the multipartite genome of *S. meliloti* [37] by *Hin*fI REC-Seq and found such hypomethylated sites on the chromosome and both megaplasmids (S1 Table).

To validate the *Hin*fI REC-Seq approach, we conducted REC-Seq (using the methylation-sensitive *MboI* restriction enzyme) on gDNA from *Escherichia coli* K12 and *V. cholerae*, as previously determined either by SMRT sequencing or REC-Seq [10, 34]. The Dam methylase introduces m6A marks at GATCs in many γ-proteobacterial genomes [4] that protect from cleavage by *Mbo*I. As known unmethylated sites in these control experiments indeed emerged with high score (S2 Table), we conclude that *Hin*fI REC-Seq is an efficient method to detect and quantitate GANTCs that escape methylation by CcrM.

### Several hypomethylated GANTCs in *C. crescentus* are MucR target sites

Having identified hypomethylated GANTCs in the *C. crescentus* genome by *Hin*fI REC-Seq, we noted that many high scoring GANTCs lie in regions that are occupied by MucR1/2 as determined by previous <u>ch</u>romatin-<u>i</u>mmuno<u>p</u>recipitation deep-<u>seq</u>uencing (ChIP-Seq) analysis [17]. Of the hits with a score higher than 100, one third lie in MucR1/2 target sequences, and the proportion is even higher (50%) in the case of the 50 top hits (Table 1 and S1 Table). To test if MucR1/2 occludes these GANTCs from methylation by CcrM, we conducted *Hin*fI-cleavage analysis of gDNA from *WT* (NA1000) and Δ*mucR1/2* double mutant by qPCR (henceforth *Hin*fI-qPCR assay) at six MucR1/2 target sites. The *CCNA_00169* promoter (henceforth P*169*) contains four GANTCs; the *CCNA_02901* promoter (P2901), the *CCNA_01149* promoter (P1149) and the *CCNA_01083* internal sequence contain two GANTCs each; the *CCNA_02830* and *CCNA_03248* promoters (P*2830* and P*3248)* carry one GANTC each (Fig 2A). A high percentage (100%) of methylation in the *Hin*fI-qPCR assay indicates that *Hin*fI cannot cleave this site because of prior methylation by CcrM, whereas a low percentage reflects efficient cleavage of the non-methylated DNA by *Hin*fI. In *WT* gDNA these six MucR1/2-target sequences are almost completely cleaved by *Hin*fI, indicating that the GANTCs are hypomethylated in the presence of MucR1/2. However, these sites are methylated and therefore not cleaved by *Hin*fI in Δ*mucR1/2* cells (Fig 2B). As control for the specificity of the *Hin*fI-qPCR assay we conducted the same analysis on sequences that are not MucR1/2 targets harbouring either i) a hypomethylated GANTC (P_*nagA*_), ii) several methylated GANTCs (P_*podJ*_) or iii) a control sequence that does not contain GANTCs (P_*xylk*_). These controls revealed a level of amplification in the *Hin*fI-qPCR assay as predicted (S1A Fig) and showed no difference between *WT* and Δ*mucR1/2* cells. Thus, only hypomethylated sequences that are bound by MucR1/2 *in vivo* are converted to methylated GANTCs in the absence of MucR1/2.

**Fig 2.**
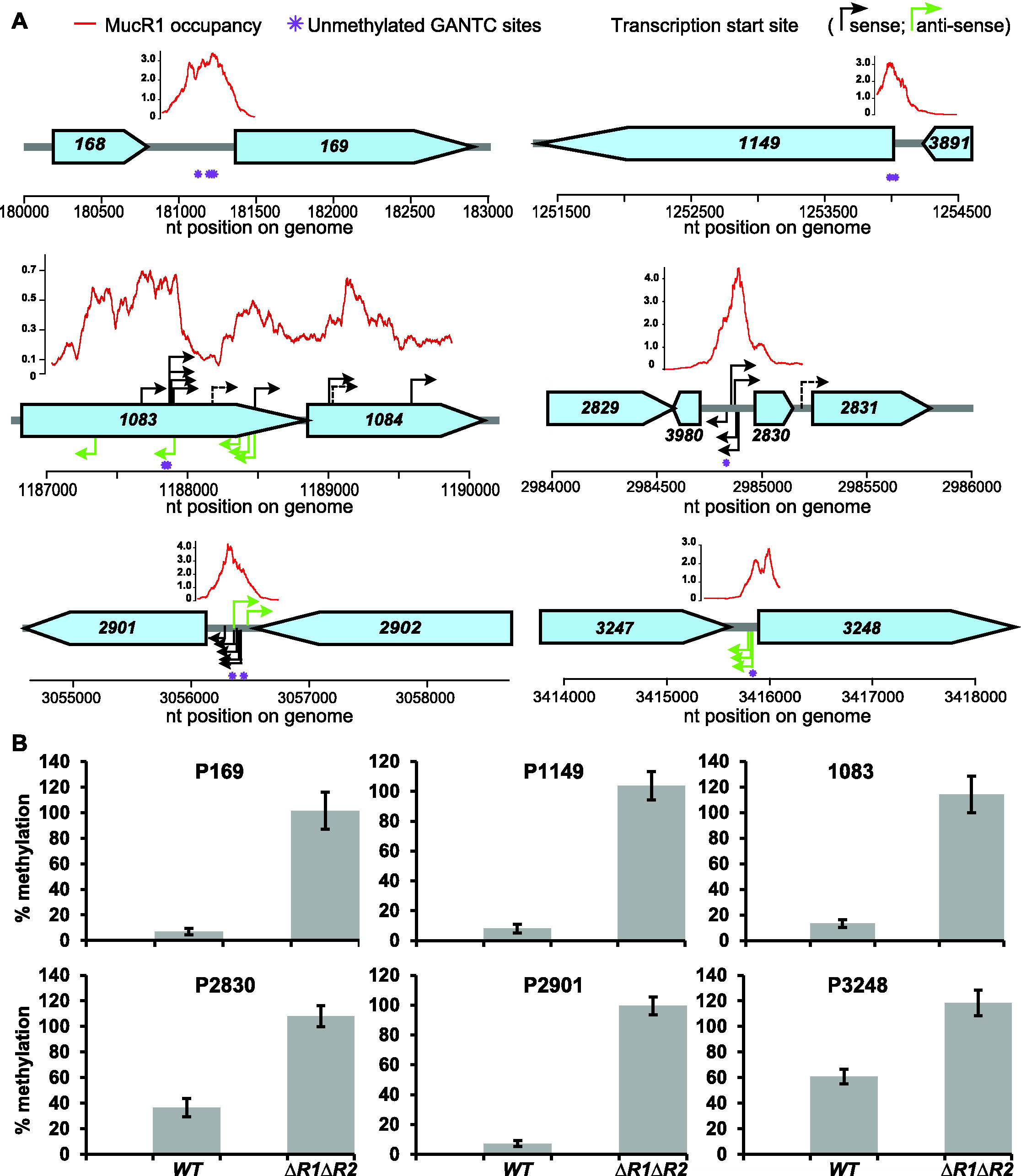
MucR occludes specific GANTC sites from methylation. (A) Schematic of the loci carrying hypomethylated GANTCs occluded by MucR. The position of the hypomethylated GANTCs identified by Kozdon et al. [12] is indicated by purple asterisks. Red lines represent the occupancy of MucR1 and the values (×10^4^ *per-base* coverage) calculated by the superresolution bioinformatic approach represent the average of the four time points (T10, T40, T70 and T100, as described in the Methods). The MucR-dependent transcription start sites, determined by TSS-EMOTE, are indicated by black (sense) and green (antisense) arrows. Dashed arrows indicate transcription start sites found in both *WT* and Δ*mucR1/2* strains (*CCNA_01083-CCNA_01084*) or down-regulated in the Δ*mucR1/2* strain compared to the *WT* (promoter of *CCNA_02831)*. (B) Methylation percentage of the loci shown in panel A in the *WT* and Δ*mucR1/2* strains, as determined by *Hin*fI-qPCR.

**Table 1.**
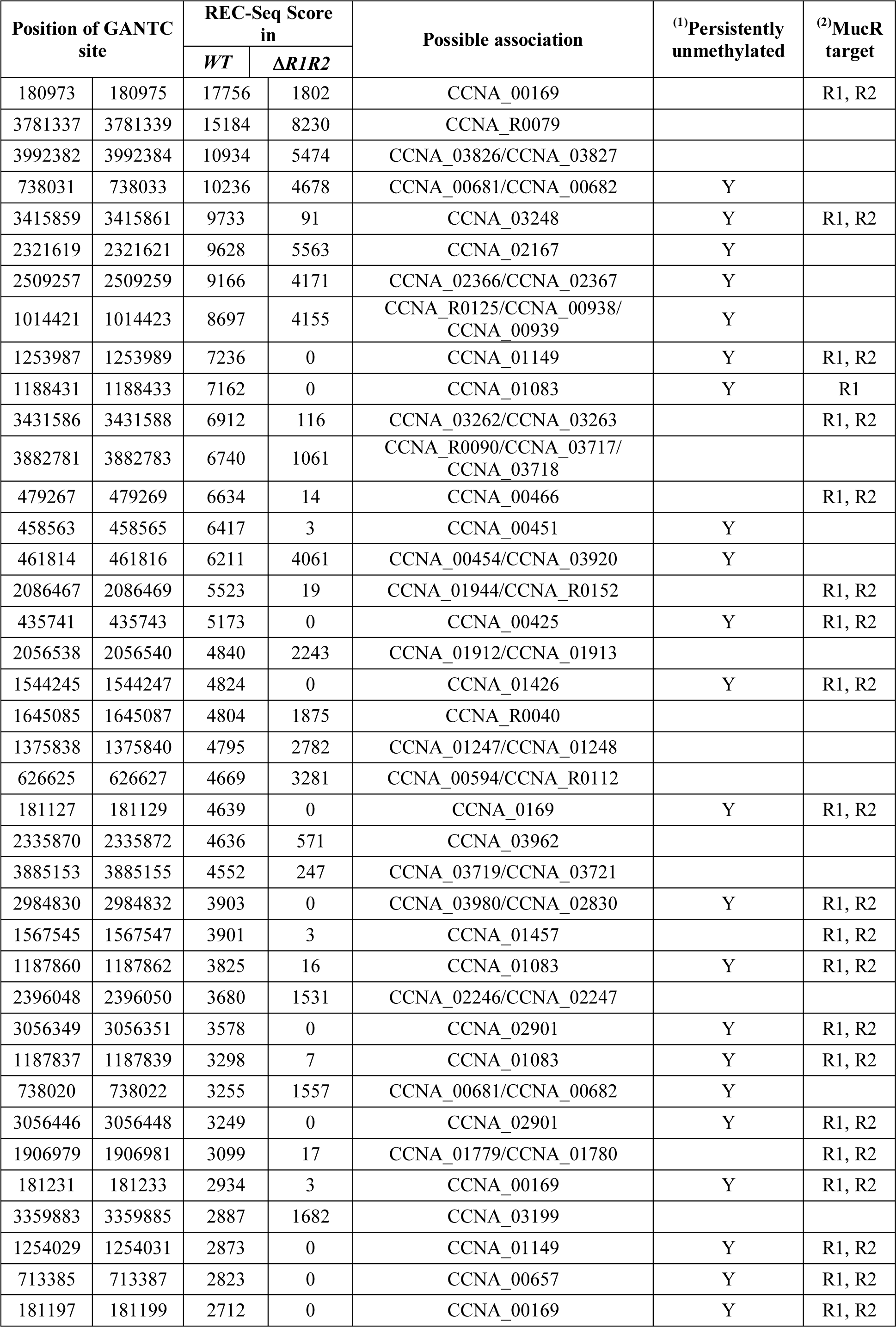
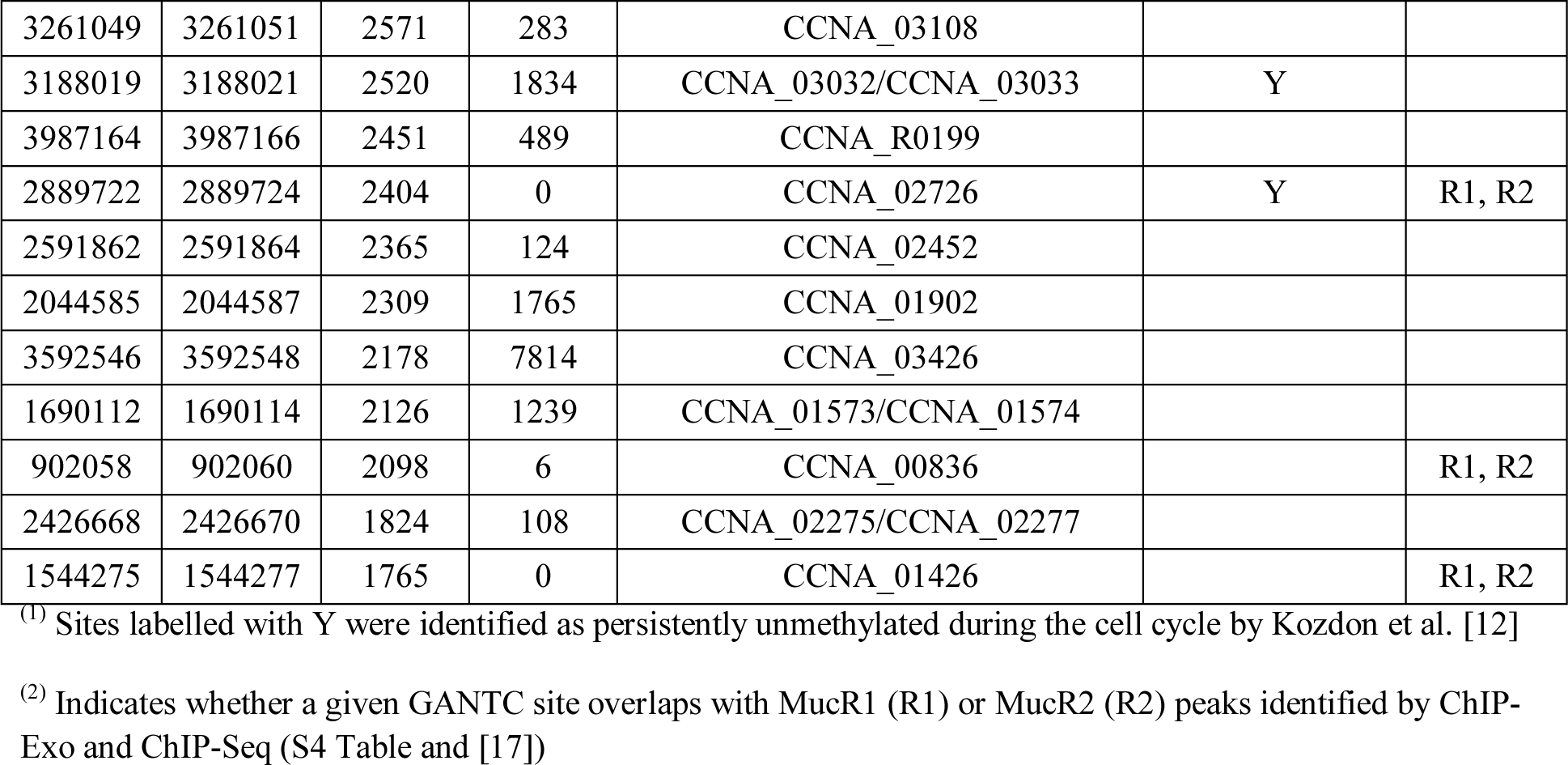
REC-Seq in *WT* and Δ*mucR1/2*. GANTC sites with the highest (top 50) REC-Seq score in *WT C. crescentus* are listed. The complete list of REC-Seq data for both *WT* and Δ*mucR1/2* strains is reported in Supplementary Table S1.

SMRT sequencing of *WT* and Δ*mucR1/2* gDNA supported the result that these GANTCs carry m6A marks as inferred by a high characteristic <u>i</u>nter<u>p</u>ulse-<u>d</u>uration (IPD) ratio observed in Δ*mucR1/2* versus *WT* cells (S3 Table). Interestingly, this analysis also revealed eleven GANTCs with the inverse behaviour, i.e. a low IPD ratio in Δ*mucR1/2* versus *WT* cells, suggesting that they no longer carry m6A marks in the absence of MucR1/2. To confirm this result we conducted *Hin*fI-qPCR assays at two of these GANTCs: the *CCNA_01248* promoter (P1248) and the *CCNA_03426* promoter *(P3426).* As predicted by the methylome analysis, we observed that the methylation percentage of these GANTCs was reduced in Δ*mucR1/2* versus *WT* (S1B Fig). On the basis of these experiments, we conclude that MucR1/2 prevents m6A-methylation by CcrM at several MucR1/2-target sequences, but can also facilitate methylation at other sites. This would likely occur by an indirect mechanism involving other MucR-dependent DNA-binding proteins that compete with CcrM at certain GANTCs.

To obtain a global picture of hypomethylated GANTCs in the absence of MucR1/2, we conducted REC-Seq analysis on gDNA extracted from the Δ*mucR1/2* strain (Table 1 and S1 Table). Comparison of the REC-Seq data for *WT* and Δ*mucR1/2* cells (S2 Fig) supported the conclusion that binding of MucR1/2 prevents methylation by CcrM, as the GANTCs tested by *Hin*fI-qPCR (shown in Fig 2) have a high REC-Seq score in *WT* and a low REC-Seq score (or they are not detected) in the Δ*mucR1/2* strain. Moreover, most of the GANTCs that show a strong decrease in score between *WT* and Δ*mucR1/2* cells are also lying in regions directly bound by MucR1/2 (Table 1 and S1 Table), based on ChIP-Exo (S4 Table) and published ChIP-Seq data [17].

### Conditions that impair local GANTC hypomethylation by MucR1/2

Since CcrM is restricted to late S-phase and MucR1/2-repression is overcome in G1-phase [17, 28], we tested if MucR1/2-bound GANTCs are still hypomethylated when CcrM no longer cycles. To this end we used two strains: the ∆*lon*::Ω (henceforth *lon*) mutant, as the Lon protease is responsible for degradation of CcrM throughout the cell cycle and upon inactivation of Lon the CcrM protein accumulates also in G1-cells, although it is only synthesized in S-phase [28, 29], and a strain with a second copy of the *ccrM* gene under control of the constitutive P_*lac*_ promoter (integrated at the *ccrM* locus, *ccrM*::P_*lac*_-*ccrM*) [30, 33]. Indeed, *Hin*fI-qPCR analysis revealed that the fraction of methylated P*169*, P*1149* and P*2901* GANTCs increases in *lon* mutant and *ccrM*::P_*lac*_-*ccrM* strain relative to *WT* cells (Fig 3A).

**Fig 3.**
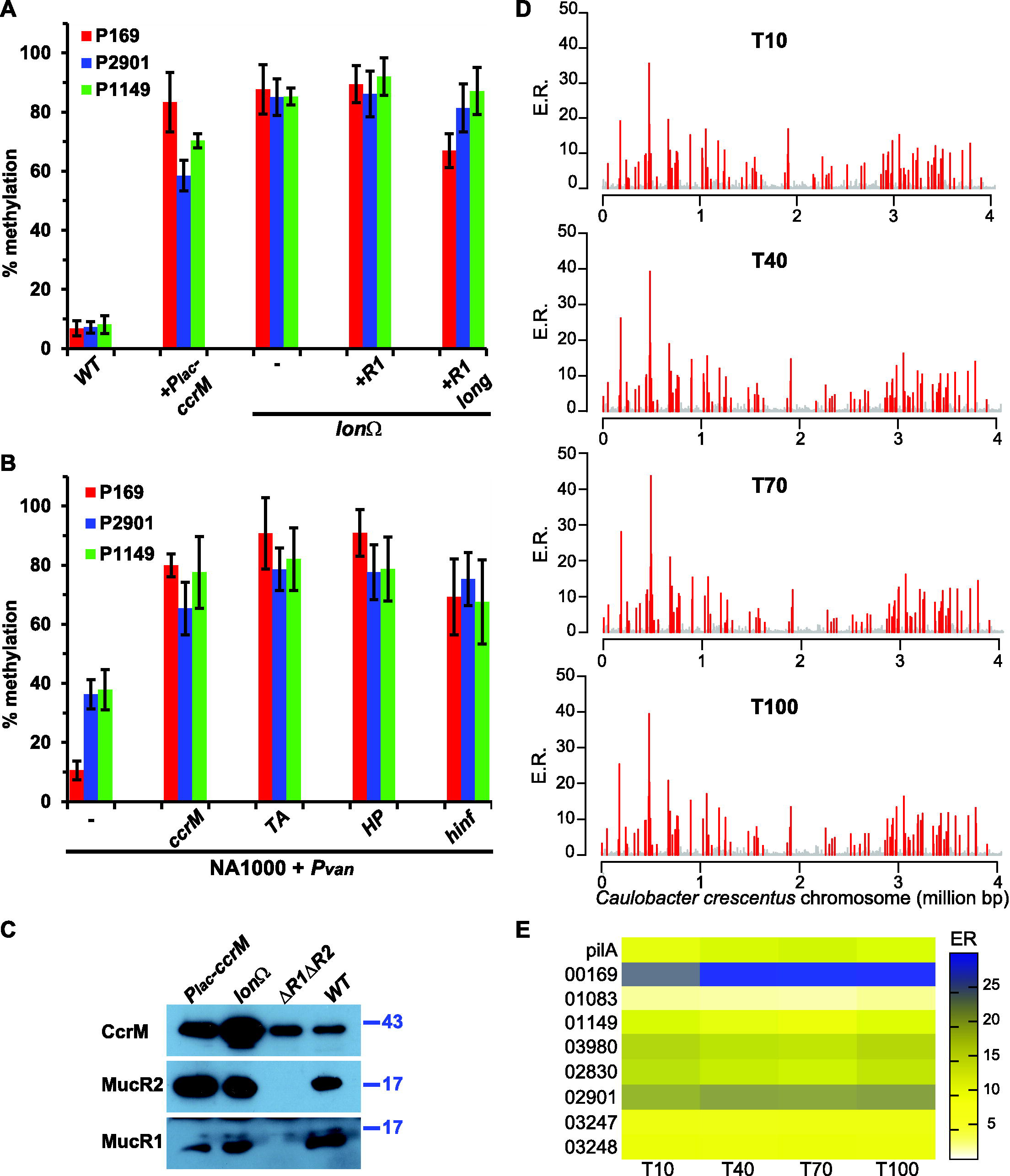
Hypomethylation by MucR is impaired in G1-phase cells. (A) Methylation percentage of the P*169*, P*1149* and P*2901* sequences in strains in which the methyltransferase CcrM is stabilised. The *Hin*fI-qPCR analysis indicates that methylation is increased in cells carrying *P*_*lac*_-*ccrM* or the *lon* mutation. In the case of the *lon* mutant, the methylation of *P169*, *P1149* and *P2901* is not affected by increased levels of MucR1 [R1: *P*_*van*_-*mucR1, R1 long:* N-terminally extended dominant-negative MucR1 variant expressed from P_*van*_ on pMT335]. (B) Methylation percentage of the *P169*, *P1149* and *P2901* sequences in *WT* cells that constitutively express *ccrM* or heterologous GANTC-methylases from P_van_ on pMT335 (TA, *Thermoplasma acidophilum;* HP, *Helicobacter pylori; hinf, Haemophilus influenzae)*. (C) Immunoblot showing the steady-state levels of CcrM, MucR1 and MucR2 in *WT* cells as well as the *P_lac_*-*ccrM, lon* and Δ*mucR1/2* strains. Molecular size standards are indicated on the right as blue lines with the corresponding values in kDa. (D) Genome-wide occupancy of MucR1 in synchronised *WT* cells at four different time points as determined by ChIP-Exo and super-resolution bioinformatic analysis: swarmer (T10), stalked (T40), early pre-divisional (T70) and late pre-divisional cells (T100). The *x* axis represents the nucleotide position on the genome, whereas the *y* axis shows the enrichment ratio (E.R.) for each promoter region as reported in the S4 Table (see Methods for a detailed description). (E) Heat map of the enrichment ratios of selected loci (those that contain hypomethylated GANTCs occluded by MucR, see Fig 2) at the four time points after synchronization. The *pilA* locus is shown as comparison as it is a well-characterized target of MucR [17]. The heat map shows that MucR1 is constitutively associated with these loci.

To exclude that constitutive presence of CcrM simply prevents MucR binding to DNA because CcrM outnumbers and therefore outcompetes MucR, we conducted several control experiments to demonstrate the specificity of the methylation control at these GANTCs. First, immunoblotting experiments revealed that MucR1/2 levels were maintained in the *lon* and *ccrM*::P_*lac*_-*ccrM* strains compared to *WT* (Fig 3C). Second, overexpression of either *WT* MucR1 or of an N-terminally extended (dominant-negative) MucR1 variant from P_*van*_ on a high copy plasmid (pMT335) [17, 38] did not prevent methylation of P169, P*1149* and P*2901* GANTCs in *lon* mutant cells (Fig 3A) or alter CcrM steady-state levels (S1C Fig). Conversely, constitutive expression of CcrM from the same vector (pMT335) in *WT* cells recapitulated the effect on methylation of the P169, P*1149* and P*2901* GANTCs (Fig 3B). Similarly, methylases of *Thermoplasma acidophilum* (TA), *Helicobacter pylori (HP*) or *Haemophilus influenzae (Hinf*), which also specifically methylate GANTCs but are not related to α-proteobacterial CcrM, also lead to methylation of these hypomethylated GANTCs when expressed from pMT335 (Fig 3B). By contrast, the methylation state of GANTCs at the *parS* locus was not significantly altered by the expression of the methylases or by the *lon* mutation (S1D Fig). On the basis that CcrM and unrelated methylases are able to compete against MucR1/2 for methylation of P*169*, P*1149* and P*2901* GANTCs when expressed constitutively, we hypothesize that MucR1/2 no longer efficiently compete with CcrM in G1-phase when both proteins are present at this time (Fig 3A, 3B).

To test if MucR1 binds to its targets in G1-phase, we conducted *ch*romatin-<u>i</u>mmunopreci<u>p</u>itation-followed by deep-<u>seq</u>uencing of <u>exo</u>nuclease treated fragments (ChIP-Exo), a technique with enhanced resolution compared to conventional ChIP-Seq [39]. We treated with the anti-MucR1 antibody chromatin prepared from synchronized cells at four different time points after synchronization [10 min (T10, G1 phase), 40 min (T40, G1-to-S transition), 70 min (T70, early S-phase), 100 min (T100, late S-phase) (Fig 1B)] and used a bioinformatic algorithm to define the binding sites at super-resolution (see Methods and [40]). Surprisingly, the binding profiles at the four time points appeared to be nearly congruent (Fig 3D) and quantification of the enrichment ratio failed to reveal major changes of MucR1 binding to its targets during the cell cycle (Fig 3E and S4 Table). On the other hand, conformational changes or altered dynamics of binding (i.e. dissociation constants, on-and off-rates) that are undetectable by our methods might allow transcription from the MucR-bound promoters in G1-phase. Transient release of DNA by MucR1/2 or changes in chromatin conformation could provide access to competing DNA binding proteins such as CcrM, RNA polymerase (RNAP) and other transcription factors (like CtrA) in G1-phase to induce methylation or firing of the MucR1/2 target promoters.

### MucR-dependent hypomethylation regulates sense and anti-sense transcription

As MucR1/2 regulates the methylation state of the aforementioned GANTCs, we wondered if the MucR1/2-targets P*169*, P*1149* and P*2901* display promoter activity in a MucR1/2-dependent and/or methylation-dependent manner. To this end, we conducted LacZ (β-galactosidase)-based promoter probe assays of P*169*-, P*1149*- and P*2901-lacZ* transcriptional reporters (driving expression of a promoterless *lacZ* gene) in *WT* and Δ*mucR1/2* cells and observed that LacZ activity of all reporters was elevated in Δ*mucR1/2* cells versus *WT* (Fig 4A-C). The increase was less dramatic for P*169*-*lacZ* (156 ± 5.8 % relative to *WT*) than for P*1149*- and P*2901-lacZ* (439 ± 7.4 and 385 ± 40 %, respectively). We then asked if promoter activity is augmented when cycling of CcrM is prevented. Indeed, the P*169*-, P*1149*- and P*2901-lacZ* reporters indicated an increase in promoter activity in the *lon* mutant and P_*lac*_-*ccrM* strains compared to *WT* (Fig 4D). Importantly, no increase in LacZ activity was observed in *lon* and P_*lac*_-*ccrM* strains with other promoters (P*hvyA* and P*pilA*, Fig 4D) that are bound by MucR1/2 and whose activity is increased in Δ*mucR1/2* cells [17, 41] but contain no hypomethylated GANTCs. We further corroborated these results by showing that constitutive expression of *C. crescentus* CcrM or the *T. acidophilum* GANTC-methylase from P_van_ on pMT335 led to an increase in P*169*-, P*1149*- and P*2901-lacZ* promoter activity (Fig 4E). Consistent with the fact that in Δ*mucR1/2* cells these promoters are no longer hypomethylated, constitutive expression of CcrM from P_*lac*_-*ccrM* in Δ*mucR1/2* cells had no significant effect on P*169*-, P*1149*- and P*2901-lacZ* promoter activity (Fig 4E).

**Fig 4.**
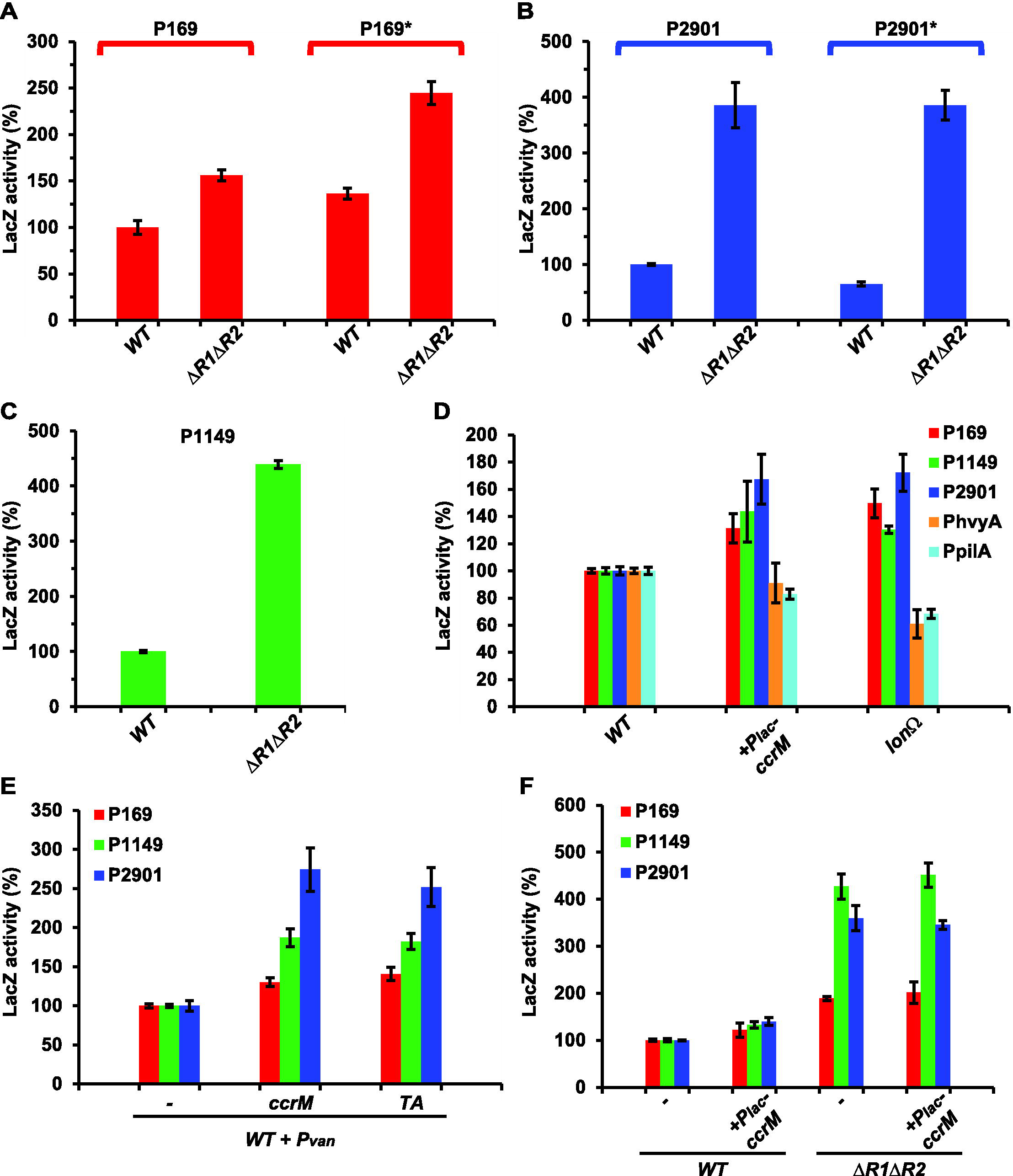
MucR and methylation by CcrM regulate transcription from target promoters. Beta-galactosidase activity of P*169* (*WT* promoter) and P*169*^*^ (with all GANTCs mutated to GTNTCs) (A), P*2901 (WT* promoter) and P*2901*^*^ (with the two GANTCs mutated to GTNTCs) (B) and P*1149* (C) in *WT* and Δ*mucR1/2* cells. Mutation of MucR1/2 increases expression from P*169*-, P*2901-* and P*1149-lacZ* independently from the presence of GANTCs. Values are expressed as percentages (activity of *WT* promoter in *WT* cells set at 100%). (D) Beta-galactosidase activity of P*169*-, P*1149*-, P*2901-lacZ* promoter probe constructs and two MucR-dependent control promoter reporters (P_*hvyA*_*-lacZ* and P_*pilA*_-*lacZ*) in *WT* and cells that constitutively express *ccrM* (ccrM::P_*lac*_-*ccrM* or ∆lon::Ω). Methylation of the target promoters by CcrM increases the LacZ activity. Values are expressed as percentages (activity in *WT* cells set at 100%). (E) Beta-galactosidase activity of P*169*-, P*1149*- and P*2901-lacZ* in *WT* cells that constitutively express *ccrM* or a heterologous GANTC-methylase from *T. acidophilum* (TA) on plasmid under control of P_*van*_. Values are expressed as percentages (activity in *WT* carrying the empty vector set at 100%). (F) Beta-galactosidase activity of P*169*-, P*1149*- and P*2901-lacZ* in *WT* and Δ*mucR1/2* cells that constitutively express *ccrM* (ccrM::P_*lac*_-*ccrM*). Values are expressed as percentages (activity in *WT* cells set at 100%).

To determine if changing the GANTC methylation state (by mutation to GTNTC) in P*169*- and P*2901-lacZ* (5 sites mutated for P*169*, P*169*^*^; 2 sites for P*2901*, P*2901*^*^) also affects promoter activity, we measured LacZ activity of the mutant promoters in *WT* and Δ*mucR1/2* cells and found that they still exhibited MucR1/2-dependency, as the P*169*^*^- and P*2901^*^-lacZ* were still strongly derepressed in the absence of MucR1/2 (Fig 4A, 4B). We also observed an increase (136% ± 6%) in activity of P*169*^*^-*lacZ* relative to P*169-lacZ* in *WT*,> while the activity of P*2901^*^-lacZ* was decreased compared to P*2901-lacZ*. The mutations may alter the target sequence for other regulator(s) in addition to the methylation properties, thereby affecting transcription directly or indirectly in a positive or negative fashion [42, 43]. For example, P*2901* is bound by the master cell cycle regulator CtrA *in vivo* and the Δ*mucR1/2* mutation is known to affect CtrA expression [17], whereas the P*169* promoter is affected by the phosphate starvation response (see below) [44, 45].

LacZ-based assays are a general and indirect measurement of promoter activity, but they do not pinpoint the transcription start sites (TSSs), thus cannot reveal the physical proximity of the TSS relative to the hypomethylated GANTCs. To correlate transcriptional regulation of MucR1/2 and hypomethylated GANTCs, we took advantage of the recently developed RNA-Seq-based strategy for <u>e</u>xact <u>m</u>apping <u>o</u>f <u>t</u>ranscriptome <u>e</u>nds (EMOTE) [46] that can also be used to map the (unprocessed) 5’ends of nascent transcripts that harbour triphosphate 5’end (5’-ppp). To this end, total RNA is first treated with XRN1 (to remove monophosphorylated 5’ends) and then 5’-ppp transcripts are treated with *E. coli* RppH, which converts the 5’ends to a monophosphorylated form that can be ligated to a bar-tagged RNA oligo using T4 RNA ligase [46] (S3 Fig). Comparative TSS-EMOTE analysis in total RNA extracted from *WT* and Δ*mucR1/2* cells unearthed several TSSs that are activated when MucR1/2 is absent (arrows in Fig 2A, S5 Table). Importantly, several of these TSSs were detected in close proximity to the GANTCs within MucR1/2 target sequences, including P*2901*, P*2830*, P*3248* and *CCNA_01083*. These results, therefore, validate the physical proximity and functional interplay between MucR1/2 and hypomethylated GANTCs. While for weak MucR1/2 target promoters the sequencing depth may have limited their detection by TSS-EMOTE, this analysis unexpectedly revealed several MucR-dependent antisense transcripts with potential regulatory roles (green arrows in Fig 2A). We validated the MucR1/2-dependency of two such antisense promoters (P*2902*_AS and P*3247*_AS) by LacZ-promoter probe assays and detected a substantial increase in activity of P*2902*_AS-*lacZ* and P*3247*_AS-*lacZ* in Δ*mucR1/2* versus *WT* cells (S1E Fig), indicating that these promoters (and the GANTCs within) are clearly MucR1/2 regulated in *C. crescentus*.

### Control of hypomethylation of MucR-target promoters in α-proteobacteria

Knowing that MucR is functionally interchangeable in α-proteobacteria [17, 18] and that hypomethylated GANTCs are also detected in the *S. meliloti* multipartite genome by *Hin*fI REC-Seq (see above and S1 Table), we tested whether *S. meliloti* MucR also occludes GANTCs from methylation by CcrM in target promoters. We compared the methylation of *WT* and mucR::Tn *S. meliloti* gDNA by *Hin*fI REC-Seq and SMRT-sequencing (S1 and S3 Table). Guided by these data sets, we validated hypomethylation of GANTCs at or near the *SMa1635* (SM2011_RS04470) and *SMa2245* (SM2011_RS06125) genes by *Hin*fl-restriction/qPCR analysis. We chose these GANTCs, located on the symbiotic megaplasmid pSymA, to take advantage of the *S. meliloti* multipartite genome and to explore if MucR-control of hypomethylation also applies to episomal elements such as a symbiotic megaplasmid. *Hin*fl-restriction/qPCR analysis revealed that these GANTCs are largely hypomethylated in *WT* compared to *mucR::Tn* cells (Fig 5A, 5B). To confirm that these GANTCs are indeed direct targets of *S. meliloti* MucR, we conducted quantitative ChIP (qChIP) experiments (Fig 5D) with chromatin from *S. meliloti WT* and *mucR::Tn* cells precipitated using antibodies to *C. crescentus* MucR2 that recognize *S. meliloti* MucR on immunoblots (S4A Fig). The qChIP experiments revealed that *S. meliloti* MucR indeed binds at or near the hypomethylated *SMa1635* and *SMa2245* GANTCs of *WT* cells (Fig 5D), but not at a control site (*SMc01552)*. Moreover, since CcrM is restricted to late S-phase also in *S. meliloti* [14], we tested whether constitutive expression of *ccr'M*^*Cc*^ in *S. meliloti WT* cells affected the methylation of GANTCs at *SMa1635* and *SMa2245*. Ectopic expression of *ccrM*^*Cc*^ from P_*lac*_ on pSRK vector [47] significantly increased the methylation of *SMa1635* and *SMa2245*, showing that *S. meliloti* MucR no longer occludes GANTCs in target promoters when cycling of CcrM is impaired (Fig 5C). Consistent with *SMa1635* and *SMa2245* being MucR targets, LacZ-based promoter probe experiments (using Pa*1635-lacZ* and Pa*2245-lacZ*) revealed that they are de-repressed in *S. meliloti mucR::Tn* cells compared to *WT* (Fig 5E) and that *S. meliloti* MucR represses Pa*1635-lacZ* and Pa*2245-lacZ* in *C. crescentus WT* or Δ*mucR1/2* cells (Fig 5F). Importantly, when cycling of CcrM in *C. crescentus* was prevented by P*_lac_*-*ccrM* or the *lon* mutation Pa*1635-lacZ* and Pa*2245-lacZ* activity was increased compared to the *WT* strain (Fig 5G). Thus, MucR controls hypomethylation during α-proteobacterial cell cycle.

**Fig 5.**
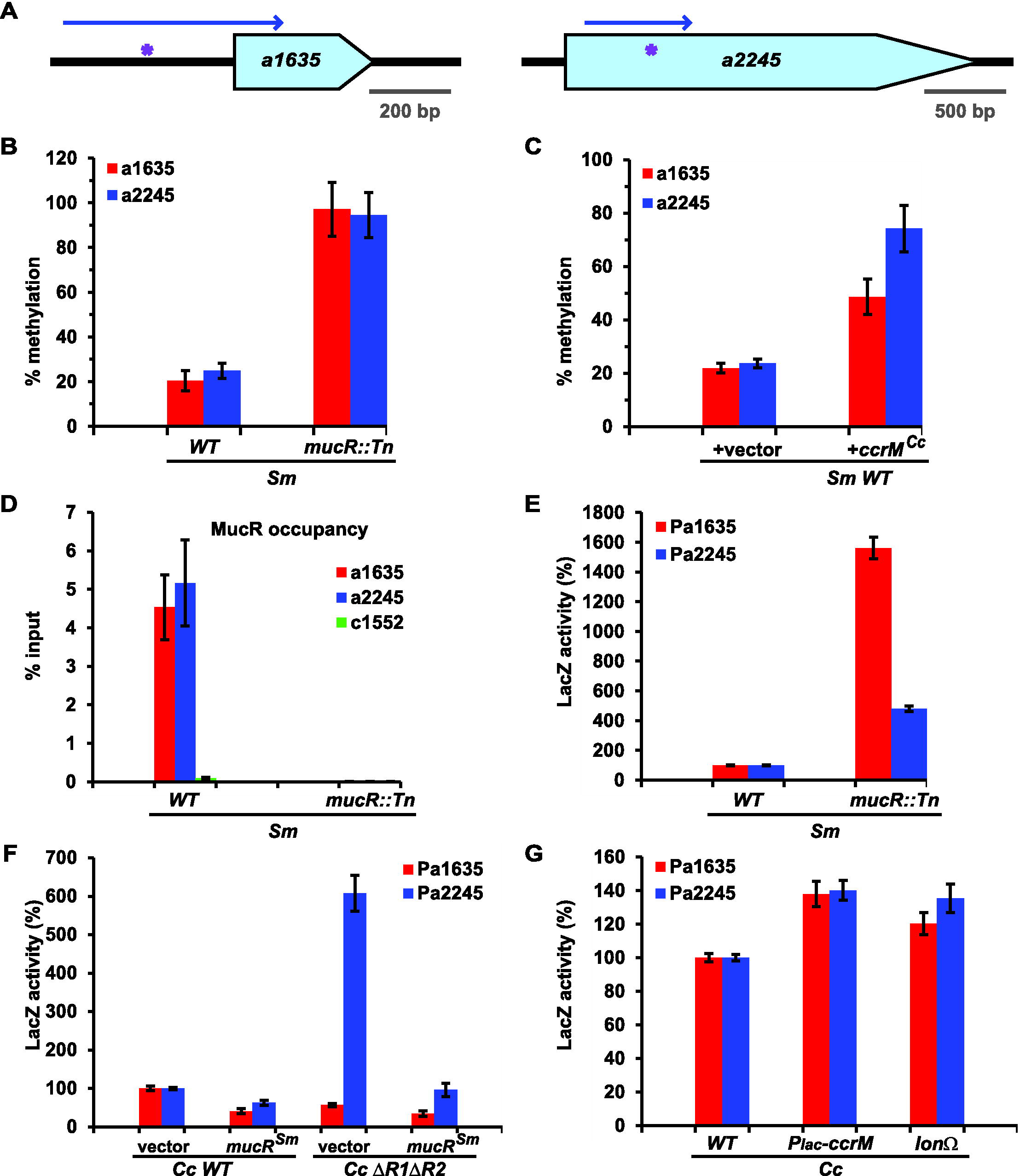
Hypomethylation control by MucR is conserved in α-proteobacteria. (A) Schematic of the two MucR-dependent hypomethylated loci (*SMa1635* and *SMa2245*) identified by SMRT-sequencing in the in *S. meliloti WT* genome. Position of the hypomethylated GANTCs is indicated by purple asterisks. The blue arrows indicate the DNA fragments cloned for LacZ promoter probe assays. (B) *Hin*fI-qPCR analysis showing that *SMa1635* and *SMa2245* are hypomethylated in *S. meliloti WT* cells compared to *mucR::Tn* cells. (C) Constitutive expression of *ccrM*^*Cc*^ from P_*lac*_ on pSRK [47] in *S. meliloti WT* cells increases the methylation percentage of *SMa1635* and *SMa2245*, indicating that hypomethylation of GANTCs by MucR is also impaired in *S. meliloti* G1-phase cells. (D) MucR occupancy at *SMa1635, SMa2245* and *SMc1552* (control) in *WT* and *mucR::Tn S. meliloti* cells, as determined by qChIP using antibodies to *C. crescentus* MucR2. *SMa1635* and *SMa2245* are bound by *S. meliloti* MucR, which suggests that hypomethylation of GANTCs at these loci is directly due to occlusion by MucR. (E) Beta-galactosidase activity of Pa*1635-lacZ* and Pa*2245-lacZ* in *S. meliloti* (fragments indicated by blue arrows in panel **A**). Both DNA fragments show a promoter activity that is strongly de-repressed in *mucR::Tn* cells compared to the *WT* strain. Values are expressed as percentages (activity in *WT* cells set at 100%). (F) Beta-galactosidase activity of Pa*1635* and Pa*2245* in *C. crescentus WT* and Δ*mucR1/2* cells expressing *mucR*^*Sm*^. Expression of *mucR*^*Sm*^ from P_*van*_ on pMT335 decreases beta-galactosidase activity of Pa*1635* and Pa*2245*. Values are expressed as percentages (activity in *WT* cells carrying the empty vector set at 100%). (G) Beta-galactosidase activity of Pa*1635* and Pa*2245* in *C. crescentus *WT*, P_*lac*_-ccrM* or *lon* cells. Values are expressed as percentages (activity in *WT* cells set at 100%).

### Environmental and systemic signals controlling hypomethylation patterns

To determine if other systemic (cell cycle) signals can alter methylation patterns in α-proteobacteria, we tested if CtrA can also occlude GANTC sites from methylation by CcrM. First, we constructed a synthetic promoter in which three GANTCs overlapping two CtrA-boxes (one GANTC in each CtrA box and one in between) were placed downstream of an attenuated *E. coli* phage T5 promoter on the *lacZ* promoter probe plasmid (Fig 6A). Next, we determined the methylation percentage of the GANTCs in *WT C. crescentus* cells harbouring the resulting reporter plasmid by *Hin*fI-qPCR analysis and found that the GANTCs are only partially methylated in *WT* cells, but efficiently methylated when CcrM is expressed ectopically (Fig 6A). Thus, methylation patterns can also emerge from competition between CcrM and other cell cycle regulators such as CtrA at appropriately positioned GANTCs.

**Fig 6.**
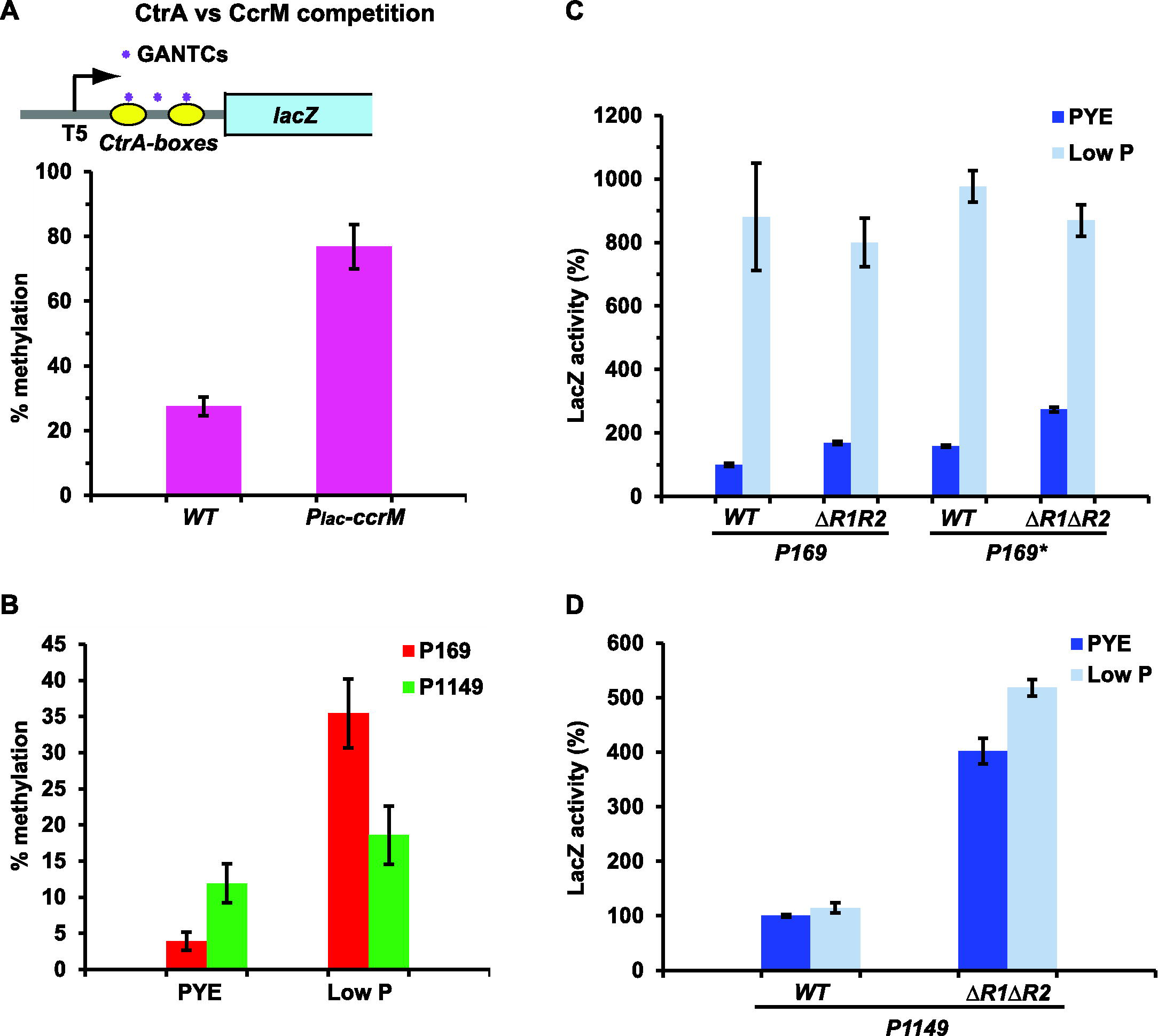
Cell cycle and environmental signals affect methylation patterns. (A) Competition between CtrA and CcrM. Schematic of the synthetic promoter carrying an attenuated *E. coli* phage T5 promoter followed by three GANTCs (purple asterisks) overlapping two CtrA-boxes (in yellow). *Hin*fI-qPCR analysis shows that this sequence is hypomethylated in *WT* cells, whereas constitutive expression of *ccrM (P_*lac*_-ccrM*) increases the methylation percentage. This indicates that DNA-binding proteins other than MucR can also occlude GANTCs from methylation. (B) Methylation percentage of P*169* and P*1149* in phosphate-limiting conditions compared to rich medium (PYE), determined by *Hin*fI-qPCR analysis. Phosphate starvation (6h) significantly increases the methylation level of P*169* but not P*1149*. (C) Beta-galactosidase activity of *P169-lacZ* and *P169*^*^-lacZ (GANTCs mutated to GTNTCs as in Fig 4A) in *WT* and Δ*mucR1/2* cells in rich medium and phosphate-limiting conditions. Phosphate starvation induces transcription from P*169-lacZ* and *P169*^*^-lacZ independently from the presence of MucR1/2. Values are expressed as percentages (activity in *WT* cells grown in PYE set at 100%). (D) Beta-galactosidase activity of P*1149-lacZ* in *WT* and Δ*mucR1/2* cells in rich medium and phosphate-limiting conditions. Phosphate starvation does not significantly affect the activity of P*1149-lacZ*. Values are expressed as percentages (activity in *WT* cells grown in PYE set at 100%).

To explore if environmental signals can also affect local hypomethylation patterns, we took advantage of the fact that expression of *CCNA_00169* (also known as *elpS*) is induced upon phosphate starvation of *C. crescentus* cells [44, 45]. Accordingly, we compared the P*169* methylation patterns by *Hin*fI-qPCR analysis of gDNA from *WT* cells grown in standard medium (PYE) and phosphate-limiting conditions. This revealed a significant increase in P*169* GANTC methylation in phosphate-limiting conditions compared to PYE (Fig 6B) and we observed a commensurate induction of P*169*-*lacZ* and P*169^*^-lacZ* that was MucR1/2 independent (Fig 6C). Both the increase in P*169* GANTC methylation and P*169*-*lacZ* activity are dependent on the phosphate-responsive transcriptional regulator PhoB (S4B-C Fig), suggesting that PhoB can facilitate methylation of MucR-protected GANTCs at P*169*. The result that no significant increase of the P*1149* methylation or P*1149-lacZ* activity was seen when *WT* cells were grown in phosphate-limiting conditions compared to standard PYE medium (or in Δ*phoB::Q*. cells compared to *WT*) argues against the possibility that changes in CcrM expression or activity underlie the modified methylation pattern of P*169* (Fig 6B, 6D; S4B-C Fig). Thus, P*169* provides an example of an environmental override for a promoter subject to local hypomethylation control by the cell cycle transcriptional circuitry.

## Discussion

The correlative changes between human genetic variability and local (hypo)methylation prompt the question if and how such patterns are regulated by the cell cycle and/or environmental cues. Taking advantage of bacterial genomes that are small enough for full-methylome analysis by cutting-edge REC-Seq and SMRT-sequencing, we show that local m6A-hypomethylation exists in two different α-proteobacterial lineages and that conserved cell cycle factors govern its establishment in both systems. While in γ-proteobacterial lineages transcriptional regulators are also known to compete with the Dam m6A methylase to occlude certain methylation sites, local hypomethylation patterning has not been explored in the context of the transcriptional circuitry controlling progression of the (α-proteo)bacterial cell cycle. In eukaryotes methylation heterogeneity involves 5-methyl-cytosines introduced at CpG dinucleotides [2], but recently m6A marks, instated by unknown mechanisms, have also been reported [6–8]. Reliable detection of methylation sites by SMRT-sequencing requires extensive (25-fold) coverage for adenine methylation and even higher coverage for cytosine methylation (250-fold coverage needed in some instances)[5]. Non-methylated sites are only reliably detected by elimination of sites on which methylation is detected, thus leaving an element of uncertainty for those sites classified as non-methylated based on the absence of the kinetic signature for methylation. By contrast, REC-Seq with a methylation sensitive restriction enzyme was used here to enrich for non-methylated sites in α-proteobacteria by *Hin*fI cleavage. The continuum of scores we detected in these experiments points towards the use of REC-Seq in detecting loci whose methylation is variable within a culture, for example phase variable loci [48, 49]. *Hin*fI REC-Seq revealed the occurrence of non-methylated GANTCs in at least two α-proteobacterial genomes. Subsequent genetic analyses showed that the determinants controlling hypomethylation are conserved in these bacteria, but they are not encoded in eukaryotic genomes. However, at least one component, MucR, possesses an ancestral zinc-finger-type DNA binding domain [22], a protein domain which is also wide-spread in developmental regulation of eukaryotes [50]. The fact that MucR regulates expression of virulence and cell cycle genes [17–20], has recently been shown to confine genetic exchange by generalized transduction to G1-phase in *C. crescentus* via transcriptional regulation [41] and is responsible for hypomethylation of specific loci on the chromosome or megaplasmids thus raises the possibility that zinc-finger proteins may control (epigenetic) DNA transactions including local hypomethylation during the eukaryotic cell cycle as well. In bacteria local methylation changes may correlate with altered virulence behaviour and may underlie cell cycle-control in pathogens, endosymbionts or other microbial systems. Methylation is known to influence virulence functions in γ-proteobacteria, often by imposing bistability from phase-variable virulence promoters in subpopulations via transcriptional regulators such as Lrp, Fur or OxyR [5, 9, 42, 48, 51–55]. Phenotypic heterogeneity in antibiotic drug tolerance *in vivo* (a state known as persistence), which is acquired in a low fraction of bacterial cells, may also underlie epigenetic changes induced stochastically by methylation, either deterministically (during the cell cycle) or environmentally. Although no phase-variable promoters are currently known for the α-proteobacteria, these bacteria offer the possibility to investigate the relationship of local hypomethylation with cell cycle control, as both *C. crescentus* and *S. meliloti* are synchronizable and exhibit comparable cell cycle control systems and transcription [13–15, 56]. However, as binding of MucR to DNA is not impaired by methylation, the mechanisms underlying the increase in transcription of target genes induced by methylation in α-proteobacteria (Fig 4D–F; Fig 5G) are likely to be different from those described for *γ*-proteobacteria. Moreover, the α-proteobacteria lineage includes the Rickettsiales order encompassing obligate intracellular pathogens, endosymbionts and the extinct proto-mitochondrion from which the modern day mitochondria descended [57]. As MucR and CcrM orthologs are not encoded in most Rickettsiales genomes, the determinants of hypomethylation in this order must be different, if they do exist. Interestingly, endosymbionts from the genus *Wolbachia* might provide a possible exception. Their genomes encode an unusual putative DNA methylase in which a C-terminal pfam01555 methylase-domain is fused to a pfam02195 ParB-like nuclease domain found in DNA-binding proteins and plasmid replication factors [58]. The sheltered niche of obligate intracellular Rickettsia contrasts with that of free-living relatives that are exposed to major environmental fluctuations.

In summary, our work shows that environmental regulatory responses like that to phosphate limitation, which is particularly pertinent for bacteria living in aquatic ecosystems as *C. crescentus*, are superimposed on (direct or indirect) hypo-or hyper-methylation control cued by the cell cycle. As many hypomethylated sites occur upstream of genes encoding transcription factors (see S1 Table) and transcription factors are often regulatory, it is conceivable that local hypomethylation is often induced by *cis*-encoded site-specific DNA-binding proteins that can compete with DNA methylases for overlapping target sites. The mechanism of DNA binding and temporal regulation of MucR remain to be elucidated in detail in order to reveal why MucR shields certain target sites from methylation by CcrM. Our work on MucR-dependent hypomethylation by *Hin*fI REC-Seq along with the comprehensive analysis of hypomethylated sites in other α-proteobacterial genomes [10] indicates that the functions controlled by hypomethylated promoters are distinct and generally not conserved among different α-proteobacteria. This suggests that hypomethylation does not play a major role in the regulation of the α-proteobacterial cell cycle, even though conserved cell cycle transcriptional regulators govern hypomethylation patterns. If it is largely serendipitous which sites MucR shields from methylation, it seems plausible that such hypomethylation control systems mediate species-specific transcriptional adaptations in response to stresses via MucR, CcrM or other variables that influence their binding, either directly or indirectly. For example, cell cycle controlled changes in local chromosomal topologies mediated by DNA replication or nucleoid-associated factors [59, 60] could exclude DNA methylases form specific target sites.

## Materials and Methods

### Strains and growth conditions

*Caulobacter crescentus* NA1000 [61] and derivatives were grown at 30°C in PYE (peptone-yeast extract) or M2G (minimal glucose). For phosphate starvation, *Caulobacter* cells were grown in 1/5X PYE (5-fold diluted PYE except 1 mM MgSO_4_ and 1 mM CaCl_2_, supplemented with 0.2% glucose). *Sinorhizobium meliloti* Rm2011 and derivatives were grown at 30°C in Luria broth (LB) supplemented with CaCl_2_ 2.5 mM and MgSO_4_ 2.5 mM. *Escherichia coli* S17-1 λ*pir* and EC100D were grown at 37°C in LB. Swarmer cell isolation, electroporations, bi-parental matings and bacteriophage ϕCr30-mediated generalized transductions were performed as previously described [62–65]. Nalidixic acid, kanamycin, gentamicin and tetracycline were used at 20 (8 for *S. meliloti*), 20, 1 (10 for *E. coli* and *S. meliloti*) and 1 (10 for *E. coli* and *S. meliloti*) µg/mL, respectively. Plasmids for β-galactosidase assays were introduced into *S. meliloti* by bi-parental mating and into *C. crescentus* by electroporation. Strains and plasmids constructions are detailed in the S1 Text file.

### Extraction of genomic DNA and methylation by qPCR (*Hin*fT-restriction/qPCR)

Genomic DNA was extracted from mid-log phase cells (10 ml). Aliquots of DNA (0.5-1 µg) were digested with *Hin*fI restriction endonuclease and used to determine the methylation percentage by Real-Time PCR. Real-time PCR was performed using a Step-One Real-Time PCR system (Applied Biosystems, Foster City, CA) using 0.05% of each DNA sample (5 µl of a dilution 1:100) digested with *Hin*fI, 12.5 µ.l of SYBR green PCR master mix (Quanta Biosciences, Gaithersburg, MD) and primers 10 each, in a total volume of 25 µl. A standard curve generated from the cycle threshold (C_t_) value of the serially diluted non-digested genomic DNA was used to calculate the methylation percentage value for each sample. Average values are from triplicate measurements done per culture, and the final data was generated from three independent cultures per strain and condition. The primers used for Real-Time PCR are listed in Table B in the S1 Text file.

### Genome-wide methylation analyses

SMRT (single-molecule real-time) sequencing libraries were prepared from gDNA extracted from the four samples (*C. crescentus* and *S. meliloti WT* and *mucR* mutant strains) using the DNA Template Prep Kit 2.0 (250bp - 3Kb, Pacific Biosciences p/n 001-540-726). Sequences generated by the Pacific Bioscience RSII were aligned to the *C. crescentus* NA1000 or *S. meliloti* Rm2011 genomes [37, 66, 67] using Blasr (https://github.com/PacificBiosciences/blasr) and the modification and associated motifs patterns were identified applying the RS_Modification_and_Motif_Analyisis protocol in SMRT Analysis (https://github.com/PacificBiosciences/SMRT-Analysis/wiki/SMRT-Analysis-Software-Installation-v2.2.0). For each aligned base, a statistics measured as interpulse duration (IPD) combined with a modification quality value (QV) would mark the methylation status. On the one hand, a minimum QV of 45 is required for a position to be marked as methylated; on the other hand, a maximum QV between 10 and 30 (depending on the observed kinetic detections background in the sample), coupled with the requirement that such a score is observed on both strands, would mean that a position, in an otherwise methylated motif, is unmethylated.

For REC-Seq (*r*estriction *e*nzyme *c*leavage-*seq*uencing) 1 µg of genomic DNA from *C. crescentus* NA1000 and *S. meliloti* Rm2011 was cleaved with *Hinf*I, a blocked (5’biotinylated) specific adaptor was ligated to the ends and the ligated fragments were then sheared to an average size of 150-400 bp (Fasteris SA, Geneva, CH). Illumina adaptors were then ligated to the sheared ends followed by deep-sequencing using a Hi-Seq Illumina sequencer, and the (50 bp single end) reads were quality controlled with FastQC (http://www.bioinformatics.babraham.ac.uk/projects/fastqc/). To remove contaminating sequences, the reads were split according to the *Hin*fI consensus motif (5’-G^A^ANTC-3’) considered as a barcode sequence using fastx_toolkit (http://hannonlab.cshl.edu/fastxtoolkit/) (fastx_barcode_splitter.pl –bcfile barcodelist.txt –bol-exact). Most of the reads (more than 90%) were rejected, and the reads kept were remapped to the reference genomes [37, 66, 67] with bwa mem [68] and samtools [69] to generate a sorted bam file. The bam file was further filtered to remove low mapping quality reads (keeping AS >= 45) and split by orientation (alignmentFlag 0 or 16) with bamtools [70]. The reads were counted at 5' positions using Bedtools [71] (bedtools genomecov -d -5). Both orientation count files were combined into a bed file at each identified 5’-GANTC-3’ motif (where reverse counts >=1 at position N+1 and forward counts >=1 at position N-1) using a home-made PERL script. The *Hin*fI positions in the bed file were associated with the closest gene using Bedtools closest [71] and the gff3 file of the reference genomes [72]. The final bed file was converted to an MS Excel sheet (S1 and S2 Tables) with a homemade script. For the *Mbo*I-based REC-Seq, the strategy was identical except that a different adaptor was used for ligation after cleavage and the *Mbo*I consensus motif (5’-^GATC-3’) was used as barcode for filtering of *V. cholerae* O1 biovar El Tor [73] and *E. coli* K12 Ec100D gDNA mapped onto the MG1655 genome [74].

### ChlP-Exo on *C. crescentus* synchronized cells

*C. crescentus WT* cells for ChIP-Exo were taken at different time points after synchronization (10, 40, 70 and 100 minutes). After cross-linking, chromatin was prepared as previously described [17]. ChIP-Exo was performed with 2 µl of polyclonal antibodies to MucR1 at Peconic LCC (http://www.peconicgenomics.com) (State College, PA), which provided standard genomic position format files (BAM) as output using the SOLiD genome sequencer (Applied Biosystems). A custom Perl script was then used to calculate the sequencing (read) coverage per base (*per-base* coverage) for each ChIP-Exo sample. Next, we computed the enrichment ratio (ER) for each promoter region. To this end, the Perl script extracted the *per-base* coverage of a 600 bp region for each ORF (from -500 to +100 from the start codon for each ORF annotated in *C. crescentus* genome) and calculated the average coverage for each of these regions. The resulting value was then normalized with respect to the coverage of all the intergenic regions. This was done (by the Perl script) by selecting all the intergenic regions in the *C. crescentus* genome, merging them and extracting the *per-base* coverage for all these intergenic regions. The coverage was averaged for windows of 600 bp, shifting each window by 100 bp, and the mean of all resulting values was computed. The ER for each promoter region was therefore calculated as the ratio between the average coverage of the promoter region divided by the mean obtained for the intergenic regions.

### Transcriptional start sites mapping by exact mapping of transcriptome ends (EMOTE)

The transcription start sites in the NA1000 *WT* and the Δ*mucR1/2* mutant were determined by TSS-EMOTE (Transcription Start Specific Exact Mapping Of Transcriptome Ends), a global assay that reveals the sequence of the 20 first nucleotides of 5’-triphosphorylated RNA in a sample based on an XRN-1 digest of transcripts lacking the 5’ triphosphate ends [46]. The TSS-EMOTE protocol and analyses were performed according to the scheme in S3 Fig and the detailed protocol described in [75]. We used a Worst-Case (i.e.) smallest difference) model to compare the number of Unique Molecular Identifiers between the two pairs of biological replicates (i.e. mutant vs. wild-type) and provide additional information about relative expression for each of the detected TSSs. The full list of detected TSSs is shown in S5 Table and TSSs at the relevant genomic loci are indicated by black (sense) and green (antisense) arrows in Fig 2A.

### β-galactosidase assays

β-galactosidase assays were performed at 30°C. Cells (50-200 µ.l) at 0D_660nm_=0.1-0.5 were lysed with chloroform and mixed with Z buffer (60 mM Na_2_HPO_4_, 40 mM NaH_2_PO_4_, 10 mM KCl and 1 mM MgSO_4_, pH 7) to a final volume of 800 µl. Two hundred µ.l of ONPG (o-nitrophenyl-γ-D-galactopyranoside, stock solution 4 mg/ml in 0.1 M potassium phosphate, pH 7) were added and the reaction timed. When a medium-yellow colour developed, the reaction was stopped by adding 400 µl of 1M Na_2_CO_3_. The OD_420_nm of the supernatant was determined and the Miller units (U) were calculated as follows: U= (0D_420nm_^*^1000)/(0D_660nm_^*^time [in min] ^*^volume of culture used [in ml]). Error was computed as standard deviation (SD) of at least three independent experiments.

### qChlP assay on *S. meliloti*

Samples for qChIP assay were prepared from mid-log phase *S. meliloti* cells as previously described [17]. Two microliters of polyclonal antibodies to MucR2 were used for the immunoprecipitation.

Real-time PCR was performed as described for *Hin*fI-restricted genomic DNA, using 0.5% of each ChIP sample (5 µl of a dilution 1:10). A standard curve generated from the cycle threshold (C_t_) value of the serially diluted chromatin input was used to calculate the percentage input value for each sample. Average values are from triplicate measurements done per culture, and the final data was generated from three independent cultures per strain. The primers used for *SMa1635* and *SMa2245* loci were the same as for the determination of the methylation percentage of these loci (Table B in S1 Text file).

### Immunoblots

For immunoblots, protein samples were separated on SDS polyacrylamide gel, transferred to polyvinylidene difluoride (PVDF) Immobilon-P membranes (Merck Millipore) and blocked in PBS (phosphate saline buffer) 0.1% Tween20 and 5% dry milk. The anti-sera were used at the following dilutions: anti-CcrM (1:10’000) [26], anti-MucR1 [17] (1:10’000), anti-MucR2 [17] (1:10’000). Protein-primary antibody complexes were visualized using horseradish peroxidase-labelled anti-rabbit antibodies and ECL detection reagents (Merck Millipore).

Plasmids, primers, synthetic fragments and strains constructions are described in the S1 Text file.

## Data Access

Deep-sequencing data are deposited in Gene Expression Omnibus database (GEO: GSE79880).

## Acknowledgments

We thank Julien Prados for help with TSS-EMOTE bioinformatics and Laurence Théraulaz for excellent technical assistance. We thank Melanie Blokesch and Yoshiharu Yamaichi for providing *V. cholerae* gDNA.

## S1 Text file

### Strains and plasmids construction

Table A. Strains and plasmids used in this study.

Table B. Oligonucleotides used in this study.

**S1 Fig. Controls for methylation percentage by *Hin*fI-qPCR and MucR-dependent antisense transcription.**

Methylation percentage of control loci in the *WT* and Δ*mucR1/2* strains, as determined by *Hin*fI-qPCR: P_*nagA*_ (hypomethylated, MucR-independent), P_*xylx*_ (no GANTCs), P_*pod*_*J* (fully methylated, MucR-independent). The controls show that MucR does not affect the methylation of sequences that are not its direct targets. (Note that as P_*xylX*_ contains no GANTCs, the label on the y-axis should not be interpreted as methylation but cleavage percentage). (B) Methylation percentage of P*1248* and P*3426* determined by *Hin*fI-qPCR analysis. The graphs show that these sequences are hypomethylated in the Δ*mucR1/2* compared to the *WT* strain, as predicted by the SMRT-sequencing and REC-Seq. This suggests that other DNA-binding proteins, directly or indirectly MucR-dependent, can also occlude GANTCs from methylation. (C) Immunoblot anti-CcrM in *WT* and *lon* mutant cells carrying an empty vector, P_*van*_*-mucR1* or P_*van*_*-mucR1* long (dominant negative form of MucR1 with an N-terminal extension). Over-expression of MucR1 does not affect the steady state levels of CcrM (arrow). Note that CcrM levels are elevated in the *lon* mutant strain as the protein is stabilized. Molecular size standards are indicated on the right as blue lines with the corresponding values in kDa. (D) Methylation percentage of the *parS* locus in *C. crescentus WT* cells carrying the empty vector, the heterologous methylases (TA, *HP, hinf*) under control of P_van_ on pMT335, P_*lac*_-*ccrM* and in the lonQ mutant. Stabilisation of CcrM or constitutive expression of heterologous methylases does not affect the methylation state of GANTCs at the *parS* locus. (E) Beta-galactosidase activity of two MucR-dependent antisense promoters identified by TSS-EMOTE. Values are expressed as percentages (activity in *WT* cells set at 100%).

**S2 Fig. Rec-Seq comparison of GANTC methylation in *WT* and Δ*mucR1/2*.**

The Rec-Seq score of each GANTC site in *WT* and Δ*mucR1/2 C. crescentus* was normalized according to the total number of reads obtained for the strains. The graph represents the difference between the normalized score obtained for the *WT* and the normalized score for the Δ*mucR1/2* strain for each GANTC site according to the position along the chromosome. Positive values (in blue) indicate hypomethylation in the *WT*, whereas negative values (in red) indicate hypomethylation in the Δ*mucR1/2* strain. The GANTCs verified by *Hin*fI-qPCR are indicated.

**S3 Fig. Flowchart of the EMOTE assay.**

RNA is shown as thin lines and DNA as thick lines, double lines represent Illumina adaptors. (A) Cellular RNAs exist as primary transcripts with a triphosphorylated 5’end (PPP, red line) and processed transcripts with either a monophosphorylated 5’end (P, dashed grey line) or a non-phosphorylated 5’-OH end (straight grey line). The asterisk indicates the ends of interest. XRN1 is used to remove 5' monophosphorylated RNA (grey dotted line) from the total RNA samples. (B) Treatment with *E. coli* RppH converts the 5' triphosphorylated end of primary transcripts to a monophosphorylated 5’end, a substrate for ligation (C, D) to the Rp6 synthetic oligonucleotide with T4 RNA ligase 1, which does not accept triphosphorylated or non-phosphorylated substrates. A mock reaction is performed at this stage in the absence of RppH to control for background (non-specific) signals. (E, F) Reverse transcription generates cDNA from both non-phosphorylated and Rp6-ligated RNA. Open arrows indicate polymerase extension. (G) Only cDNA from Rp6-ligated (and therefore originally triphosphorylated) RNA is amplified by the primers that add EMOTE barcodes (xxx) and Illumina adaptors. (H) Illumina sequencing (50 nucleotides) from the “A” end (see panel G) results in reads that have the specific EMOTE barcode of the original RNA sample, the Rp6 sequence and the first 20 nucleotides of the original triphosphorylated RNA, permitting exact identification of the original 5' end.

**S4 Fig. Antibodies against *C. crescentus* MucR2 specifically recognize *S. meliloti* MucR. Methylation and induction of P*169* are PhoB-dependent.**

(A) Immunoblot on *C. crescentus* and *S. meliloti* total cell extracts showing that the polyclonal antibodies against *C. crescentus* MucR2 specifically recognize *S. meliloti* MucR (SMc00058). Molecular size standards are indicated on the right as blue lines with the corresponding values in kDa. (B) Methylation percentage of P*169* and P*1149* in phosphate-limiting conditions compared to rich medium (PYE) in *WT* and Δ*phoB::Q* cells, determined by *Hinf*I-qPCR analysis. The graphs show that the increase in the methylation state of P*169* is dependent on the presence of the conserved transcriptional regulator PhoB. (C) Beta-galactosidase activity of P*169-lacZ* and P*1149-lacZ* in *WT* and Δ*phoB:: Q* cells in rich medium and phosphate-limiting conditions. The induction of P*169-lacZ* is specifically dependent on the presence of PhoB. Values are expressed as percentages (activity in *WT* cells grown in PYE set at 100%).

**S1 Table**. REC-Seq *Hin*fI analysis of *C. crescentus* and *S. meliloti* genomic DNA. In both cases, wild type and *mucR* mutant strains were analysed. For the GANTC sites with a score higher than 100 in the *WT C. crescentus*, REC-Seq data were compared to available ChIP-Seq [17] and SMRT-Sequencing data [12]. The GANTCs tested by *Hin*fI-qPCR assay are highlighted in yellow. The 50 GANTCs with the highest score in *WT C. crescentus* are those shown also in Table 1.

**S2 Table**. REC-Seq *MboI* analysis of *E. coli K12* and *Vibrio* genomic DNA.

**S3 Table**. Non-methylated GANTCs predicted from SMRT analysis of *WT* and *mucR* mutant in *C. crescentus* and *S. meliloti* genomic DNA.

**S4 Table**. ChIP-Exo analysis of MucR1 occupancy at different time points (T10, T40, T70, T100) during the *C. crescentus* cell cycle.

**S5 Table**. TSS-EMOTE analysis of transcription start sites in *WT* and Δ*mucR1/2 C. crescentus* cells.

